# Protocol for generating 3D mesenchymal stromal cell spheroids using serum-free medium and differentiating them into endothelial-like Cells

**DOI:** 10.64898/2026.11.18.767559

**Authors:** Simeng Li, Isabel Arias Quiros, Owen Smith, Walter Kolch, Guenther Eissner

**Author notes:** Technical contact: Simeng Li. Lead contact: Guenther Eissner.

## Abstract

Three-dimensional (3D) cultures of mesenchymal stromal cells (MSC) replicate in vivo conditions more accurately than two-dimensional (2D) cultures. This protocol outlines a simple and cost-efficient approach for generating mesenchymal stromal cell (MSC) spheroids, which typically form within 1-3 days and are suitable for endothelial differentiation. Accutase can be applied to dissociate the spheroids into single cells for downstream analysis. A combination of growth factors is used to induce the differentiation of MSC spheroids into endothelial-like structures, which express key endothelial markers such as von Willebrand factor (vWF) and CD31, as confirmed by immunofluorescence.

**Graphical abstract:** 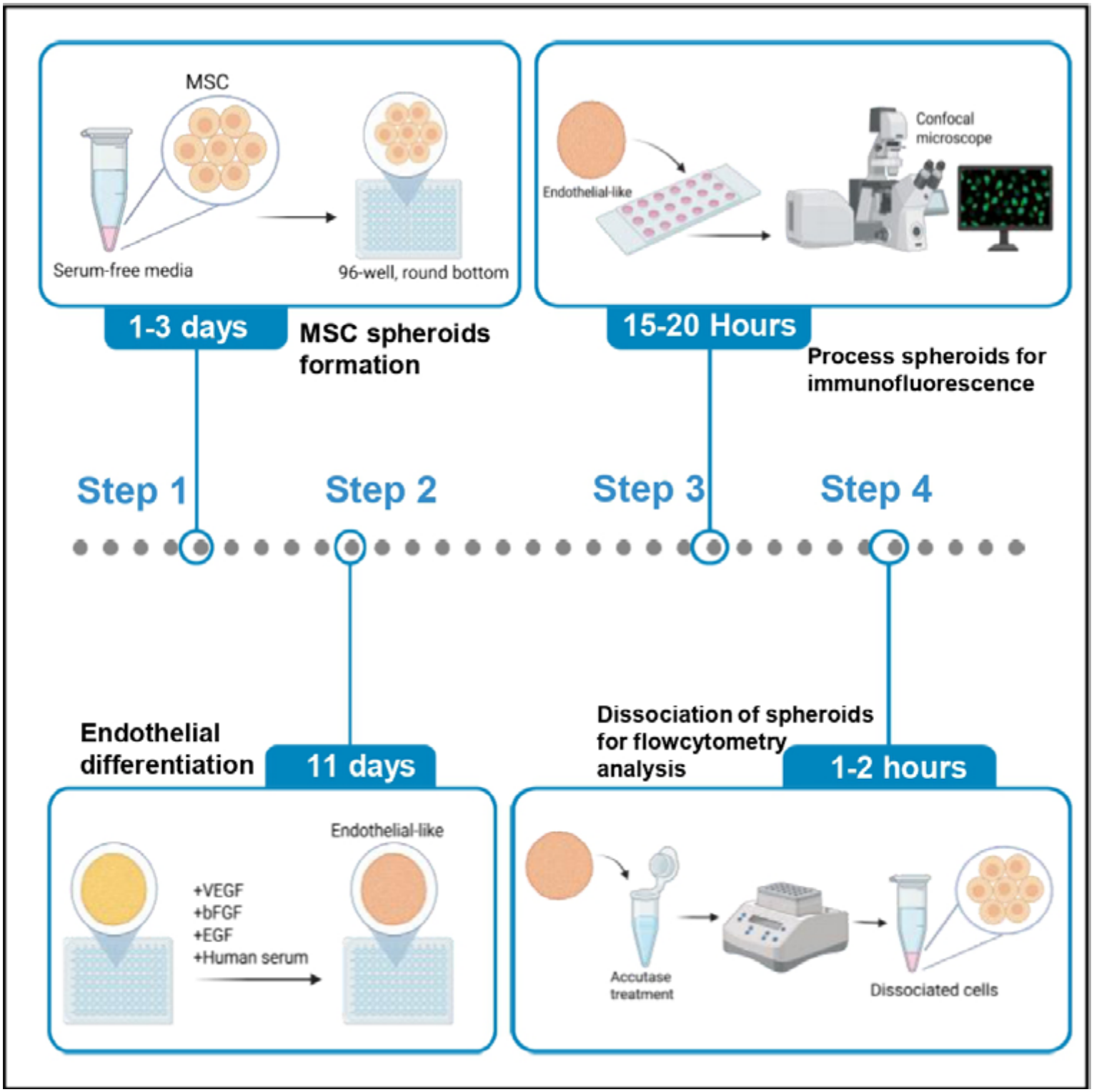

## Before you begin

Three-dimensional (3D) culture techniques have become essential in advancing MSC research, particularly for applications such as endothelial differentiation. Numerous methods for generating MSC spheroids have been developed, including hanging drop, magnetic levitation, suspension cultures, and hydrogels ^1, 2^. While these methods provide physiologically relevant models, many rely on complex reagents or equipment, making them both cost-inefficient and impractical for clinical applications. Additionally, many techniques utilize foetal bovine serum (FBS), which cannot be employed in clinical settings due to the risk of contamination and immune reactions.

Recent advancements have introduced simpler and more cost-effective methods for forming MSC spheroids in animal component free culture. For instance, commercially available serum-free or chemically defined xeno-free media, supplemented with human serum albumin (HSA), or using forced aggregation techniques in low-binding plates, allow spheroid formation in under three days ^3, 4^. Culturing MSC in serum-free Dulbecco’s Modified Eagle’s Medium (DMEM) results in the spontaneous formation of multicellular aggregates, termed sphere cluster formations (SCF), which serve as precursors to fully formed spheroids ^5^. Additionally, using low-binding 96-well plates or knockout serum replacement (KSR), a defined FBS-free formulation, enables MSC spheroids to form while retaining their key properties ^6, 7^. These advancements offer a promising, efficient pathway for generating clinical-grade MSC spheroids.

From our studies, MSC differentiated in 2D xeno-free culture was able to form tubes on Matrigel, express vWF proteins and uptake acetylated LDL. However, despite strong stimuli (hypoxia, shear stress) or extended differentiation durations applied, inducing endothelial surface markers on differentiated MSC cultured in 2D systems often falls short as observed in our studies, which did not align with other published research findings ^8-10^. In contrast, MSCs cultured in 3D spheroids exhibit significantly enhanced differentiation into endothelial-like cells ^11, 12^.

Moreover, cells derived from 3D spheroids express higher levels of endothelial markers, such as CD31 and VEGFR2, compared to those from 2D cultures^11^. The chemokine receptor CXCR4, critical in angiogenesis and MSC migration, is also upregulated in 3D spheroids ^13^. This shift in cellular marker expression, including downregulation of traditional MSC markers (CD90 and CD105) and upregulation of CD34, indicates the enhanced angiogenic potential of MSC cultured in 3D environments^12^. These features make 3D MSC spheroids a valuable tool for vascular regenerative therapies.

Finally, MSC can be recovered from 3D spheroids, retaining their original MSC characteristics, likewise, endothelial-differentiated spheroids maintain their endothelial properties upon recovery ^7, 11^. When transferred back to FBS-containing cultures, MSC migrate out of the spheroids and adhere to the plastic surface within 24 hours, although it may take up to six days for complete migration and spheroid dissolution^7, 11^. Proteolytic or collagenolytic enzymes such as trypsin, Accutase, or collagenase can also be used to dissociate the spheroids more rapidly ^14-16^. This enzymatic method enables immediate analysis via techniques like flow cytometry, allowing for assessments of cell viability, apoptosis, and proliferation without requiring additional 2D culturing steps.

This protocol, therefore, presents a straightforward and cost-efficient approach for generating, differentiating, and recovering MSC from 3D MSC spheroids, with potential applications in regenerative medicine and clinical-grade stem cell therapies.

The protocol below describes the specific steps for using hTERT20-bone marrow (BM)MSC cells.

### Preparation of coated culturewares for serum-free MSC culture

**Timing: 2 hours or overnight**

1. Flask coating prior to cell seeding
  a. Dilute the Animal Component-Free (ACF) Cell Attachment Substrate (included in the MesenCult™-ACF Plus Culture Kit) 1:150 in D-PBS (without Ca++ and Mg++)
  b. Add diluted ACF cell attachment substrate to coat the culturewares for 2 hours at Room temperature (15-25 °C)
  **Note**: using sufficient volume to cover the culture surface (around 700 µL per 10 cm2). **Note**: Alternatively, coat flasks Overnight at 2-8 °C with flask lid sealed by parafilm.
2. Remove attachment substrate completely by tilting the flask and gathering the substrate to the edge of the flask.
3. Wash flasks once with D-PBS. Then the flasks are ready to use.

Note: add DPBS to the side of the flask instead of directly pipetting onto the coated surface.

### Preparation of fully-supplemented MesenCult-ACF Plus Medium

**Timing: 20 min**

4. Complete 500 mL MesenCult-ACF Plus Medium by adding 1 mL of MesenCult-ACF Plus 500X Supplement and 2 millimolar (mM) of L-glutamine. This medium is used as normal culture medium.

### Preparation of endothelial differentiation medium

**Timing: 10 min**

5. Make endothelial differentiation medium by adding 10ng/ml (final concentration) VEGF, EGF, bFGF and 2% (v/v) human serum (HS) into fully-supplemented MesenCult ACF-plus medium

### Preparation of buffers for Immunofluorescence

**Timing: 30 min**

6. Prepare 4% of Paraformaldehyde (PFA), dilute 16% PFA at 1:4 ratio in dH_2_O to reach 4% final concentration of PFA.
7. Make blocking buffer use PBS containing the following components:
8. Prepare PBST by diluting Tween®20 detergent in PBS to reach 0.05% final concentration (v/v).
9. Prepare TBST by diluting Tween®20 detergent in Tris-buffered saline to reach 0.1% final concentration (v/v).

| Components | Concentration (v/v) |
|---|---|
| FBS | 3% |
| BSA | 1% |
| Triton X-100 | 0.50% |
| Tween<sup>®</sup>20 | 0.50% |

Note: See detailed recipe in Materials and Equipment setup section.

### Preparation of buffers for flow cytometry

**Timing: 10 min**

10. Prepare FACS buffer by adding 0.5% BSA and 2 mM EDTA into PBS. Sterile filter through 0.22 µm filter is recommended. **Note**: Buffer can be stored in 2-8 °C for 6 months. **Note:** See detailed recipe in Materials and Equipment setup section.

### Key resources table

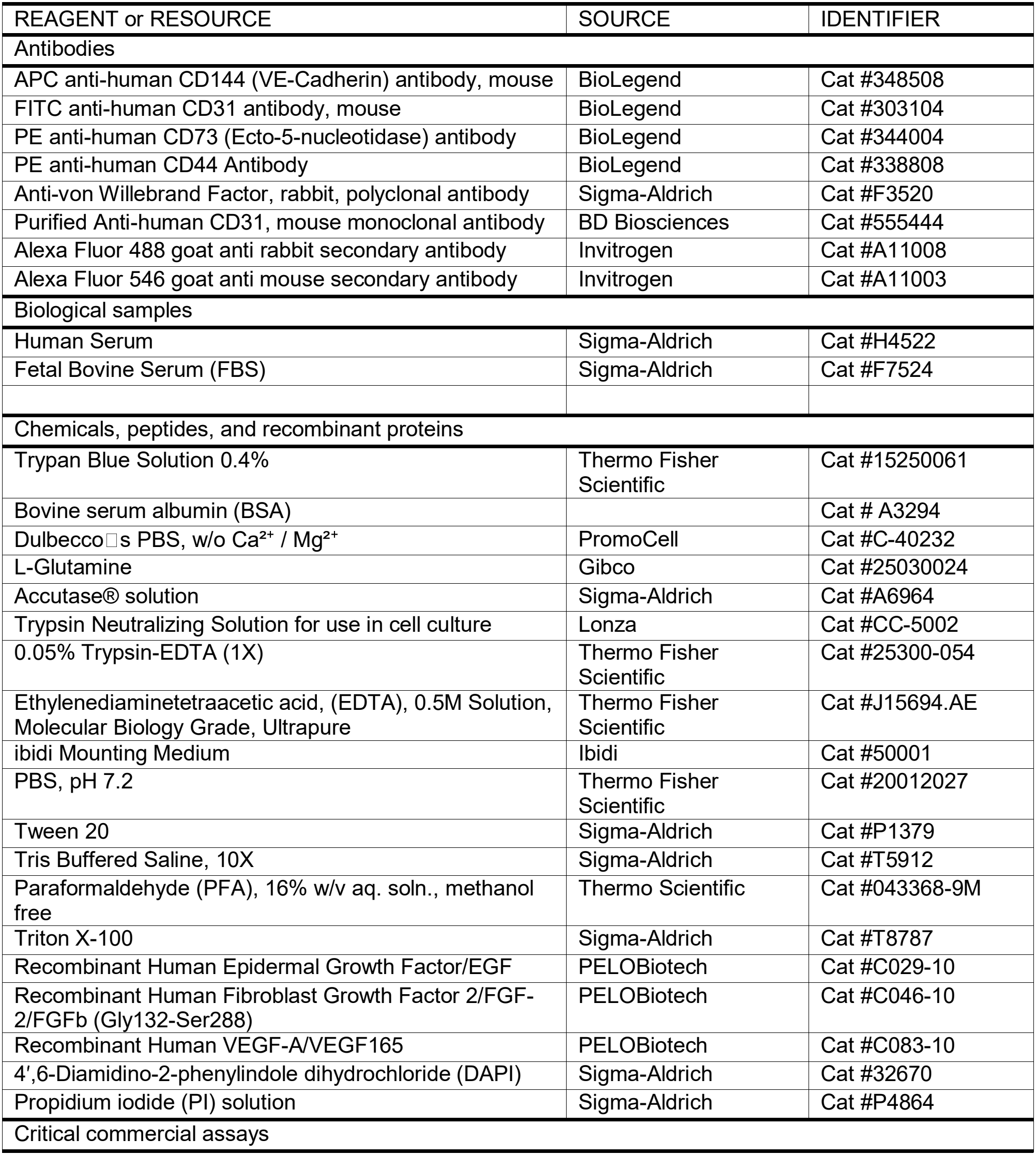
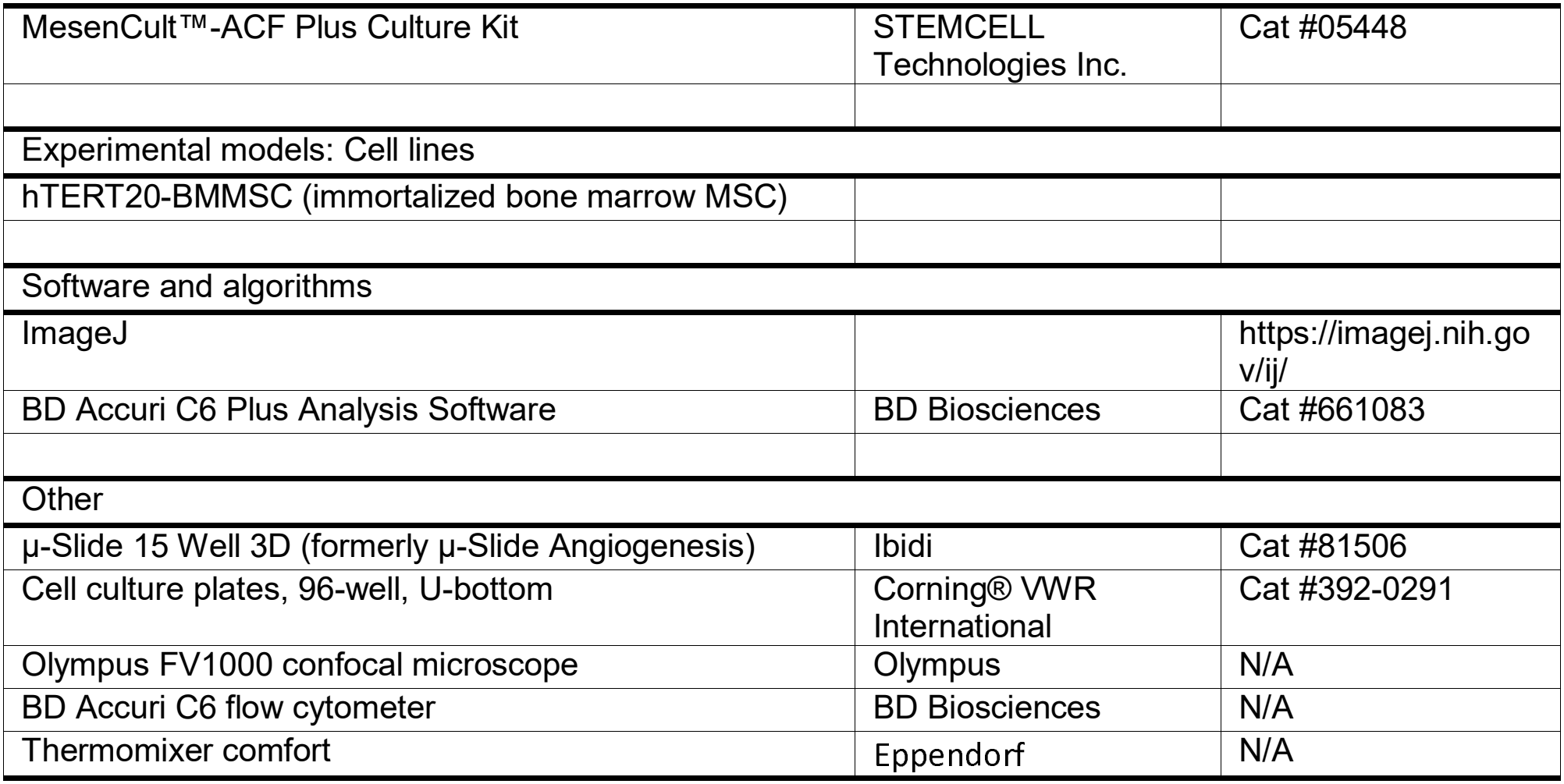

## Materials and equipment setup

### Immunofluorescence blocking buffer

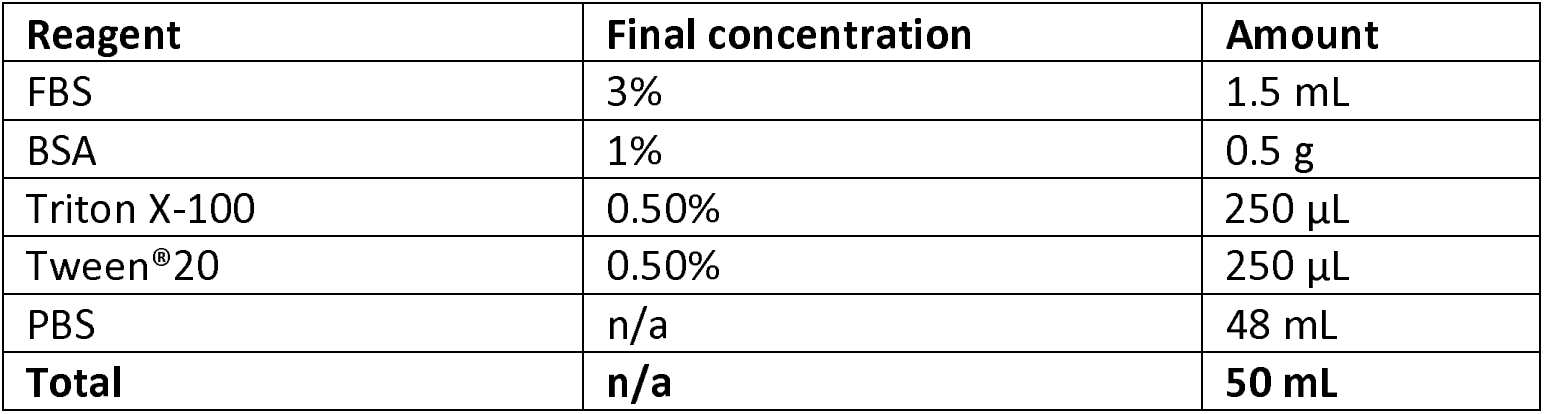

### Reagent Final concentration

| Reagent | Final concentration | Amount |
|---|---|---|
| BSA | 0.5% | 0.25 g |
| EDTA (0.5M stock solution) | 2mM | 200 µL |
| PBS | n/a | 49.8 mL |
| **Total** | **n/a** | **50 mL** |

### Step-by-step method details

#### Mesenchymal stromal cell culture and spheroids formation

**Timing: 2.5-3 hours**

This step involves trypsinizing MSCs to assess cell viability and perform cell counting. MSC with viability greater than 95% are then diluted to a specified concentration and seeded into a 96-well round-bottom plate for spheroid formation.

1. Trypsinize MSC with controlled treatment time. Normally 30 seconds to 1 minute is sufficient to detach MSC from flasks. **Note:** Use 1 mL of 0.05% trypsin-EDTA solution per 10 cm^2^. **Critical:** Use trypsin neutralizing solution to neutralize trypsin since there is no serum in the cell culture medium, 1 mL of trypsin neutralizing solution is needed to neutralize 1 mL trypsin.
2. Centrifuge neutralized cell suspension at 300 x g for 5 minutes.
3. Remove supernatant after centrifugation and resuspend the cells in 1 mL of fully-supplemented MesenCult-ACF Plus Medium.
4. Count cells with Trypan Blue exclusion using a haemocytometer and seed MSC in fully-supplemented MesenCult-ACF Plus Medium on a pre-coated cultureware. A sub-cultivation ratio of 1:3 is used. Culture condition is 37 °C, 5% CO_2_ and 80% humidity.
5. Dilute cells in fresh medium to a concentration of 3×10e5/ml. **Critical:** Cells need to be over 95% viable to form spheroids.
6. Seed 50 µL of cell suspension to each well of a 96-well round-bottom plate. There will be 15,000 cells seeded in each well.
7. Keep MSC in incubator for 1-3 days until one spheroid is formed in each well. Depending on the size of the spheroids, they can be observed either with the naked eye or with a microscope. Culture condition is 37 °C, 5% CO_2_ and 80% humidity. **Note:** multiple spheroids can be observed in each well if higher number of cells were seeded.

### Endothelial differentiation of MSC spheroids

**Timing: 11 days**

This step is for initiating spheroids differentiation after spheroids formation.

8. Carefully remove the media from each well, avoid the spheroid stucking at the end of the tips. **Note**: use a 200ul pipette tip with smaller opening
9. Add 50 µL of differentiation medium into each well. Keep some spheroids as untreated control by adding normal culture medium.
10. Replace the medium every day. Differentiation duration is 11 days.

Note: Differentiation can be determined by how advanced the differentiated cells need to be.

### Immunofluorescence to check endothelial markers of differentiated spheroids

**Timing: 15-20 hours**

This major step listed the protocol for performing immunofluorescence on the spheroids which were formed and differentiated from the two previous steps. The protocol in this section was adapted from Wimmer et al^17^.

11. Remove medium in each well carefully. Before fixation and permeabilization, wash spheroids three times with PBS, 5 min each. **Note:** From this step, the experiment doesn’t have to be performed in a sterile environment.
  a. Washing spheroids with 100 µL PBS, leave the PBS for 5 minutes
  b. Carefully remove the PBS and add fresh 100 µL PBS, leave for 5 minutes
  c. Carefully remove the PBS and add fresh 100 µL PBS, leave for 5 minutes and then discard the supernatant.
12. Use the same 96-well round-bottom plate for fixation. Perform fixation by adding 200 µL of 4% PFA into each well for 1 hour at room temperature. **Note:** The fixation time depends on spheroid size, with 1 hour being appropriate for spheroids with a diameter of less than 550 µm.
13. Wash spheroids with 100uL PBS per well three times, 5 min each. Same as Step 11.
14. Block spheroids using blocking buffer for 2 hours at room temperature on a rocker with speed setting at 35 rpm.
15. Dilute primary antibodies 1:100–1:200 (or according to manufacturer’s instruction) in the blocking buffer.
16. Remove blocking buffer from each well and add 50 µL of primary antibody to each well and incubate overnight at 2 °C – 8 °C. **Note:** primary antibodies from different species can be used together.
17. Remove solution from each well and wash spheroids with PBST for 10 min at room temperature on a rocker, wash three times.
  a. Washing spheroids with 200 µL PBST, leave for 10 minutes
  b. Carefully remove the PBST and add fresh 200 µL PBST, leave for 10 minutes
  c. Carefully remove the PBST and add fresh 200 µL PBST, leave for 10 minutes and then discard the supernatant.
18. Add corresponding secondary antibodies diluted 1:250 in blocking buffer (or according to manufacturer’s instruction), incubate for 2 hours at room temperature on a rocker.
19. Remove solution from each well and wash spheroids with TBST (0.1% Tween® detergent in Tris-buffered saline) for 20 min at room temperature on a rocker, wash three times.
  a. Washing spheroids with 200 µL TBST, leave for 20 minutes
  b. Carefully remove the TBST and add fresh 200 µL TBST, leave for 20 minutes
  c. Carefully remove the TBST and add fresh 200 µL TBST, leave for 20 minutes and then discard the supernatant.
20. Dilute DAPI stock in PBS to reach 300 nanomolar (nM) working concentration.
21. Stain Spheroids with 300 nanomolar (nM) DAPI for 30 min at 37 °C
22. Washing step. Same as step 17.
23. Use a 1 ml pipette to take the spheroid up together with the solution from each well. Transfer the spheroid to a well of ibidi µ-Slide 15 Well 3D.
24. Remove PBST from each well of ibidi µ-Slide, add a drop of ibidi mounting medium to mount the spheroids. Control the amount of mounting medium added, so that the spheroids will be located at the bottom of each well. Note: High volume of mounting medium will make spheroids float in the well and make it difficult to image.
25. Image the spheroids with an Olympus FV1000 confocal microscope (Olympus Lifescience), or other suitable confocal microscopes.

### Disassociation of MSC from Spheroids for flow cytometry analysis

**Timing: 1-2 hours**

This step uses Accutase solution for dissociating cells from spheroids for further flow cytometry analysis.

26. Harvest 10 spheroids from each condition into one 1.5 mL Eppendorf tubes. Centrifuge at 300 x g for 4 min. Remove and discard the supernatant. **Note:** use 1 mL pipette which allows spheroids been taking up with the culture medium.
27. Add 400 µL accutase solution to each tube and incubate the tube on a Thermomixer comfort at 1400 rpm, 37°C for 10 min.
28. Use a 200uL pipette to pipette up and down for ten times to further disassociate the spheroids. This is considered as one disassociation cycle.
29. Remove used Accutase solution after centrifugation at 300 x g for 4 min.
30. Repeat the disassociation cycle (step 27-29) 1-3 times more with fresh accutase each time. until no visible cell clumps remaining in the tube.
31. Wash cells once with 1 mL PBS. Centrifuge at 300 x g for 5 min and then cells can be resuspended in 100 µL FACS buffer (PBS with 0.5% BSA and 2 mM EDTA) for antibody labelling.
32. Add 5 µL of each antibody into 100 µL of cell suspension. Mix well and incubate at 2-8 °C for 30 min. Note: Adjust dilution and incubation time according to the manufacturer’s instructions for different antibodies.
33. After incubation, add 1 mL PBS to cell suspension, wash at 300 x g for 5 min. Discard supernatant.
34. Repeat step 33 two more times for a total 3 time of wash
35. Resuspend cell pellet in 100 µL FACS buffer. Cell suspension is now ready for flow cytometry analysis.
36. PI or DRAQ7 staining is essential for live/dead cell extinguish.
  a. 1 µL of PI solution was added to 100 µL cell suspension right before analysis to reach final concentration of 10 µg/ml.
  b. 1 µL of DRAQ7 solution was added to 100 µL cell suspension to reach final concentration of 3µM, incubate 10 min before analysis.
37. Run labelled samples on BD Accuri C6 flow cytometer, gate strategy is shown below: singlets - > scatter cells -> live cells-> positive population for each marker.

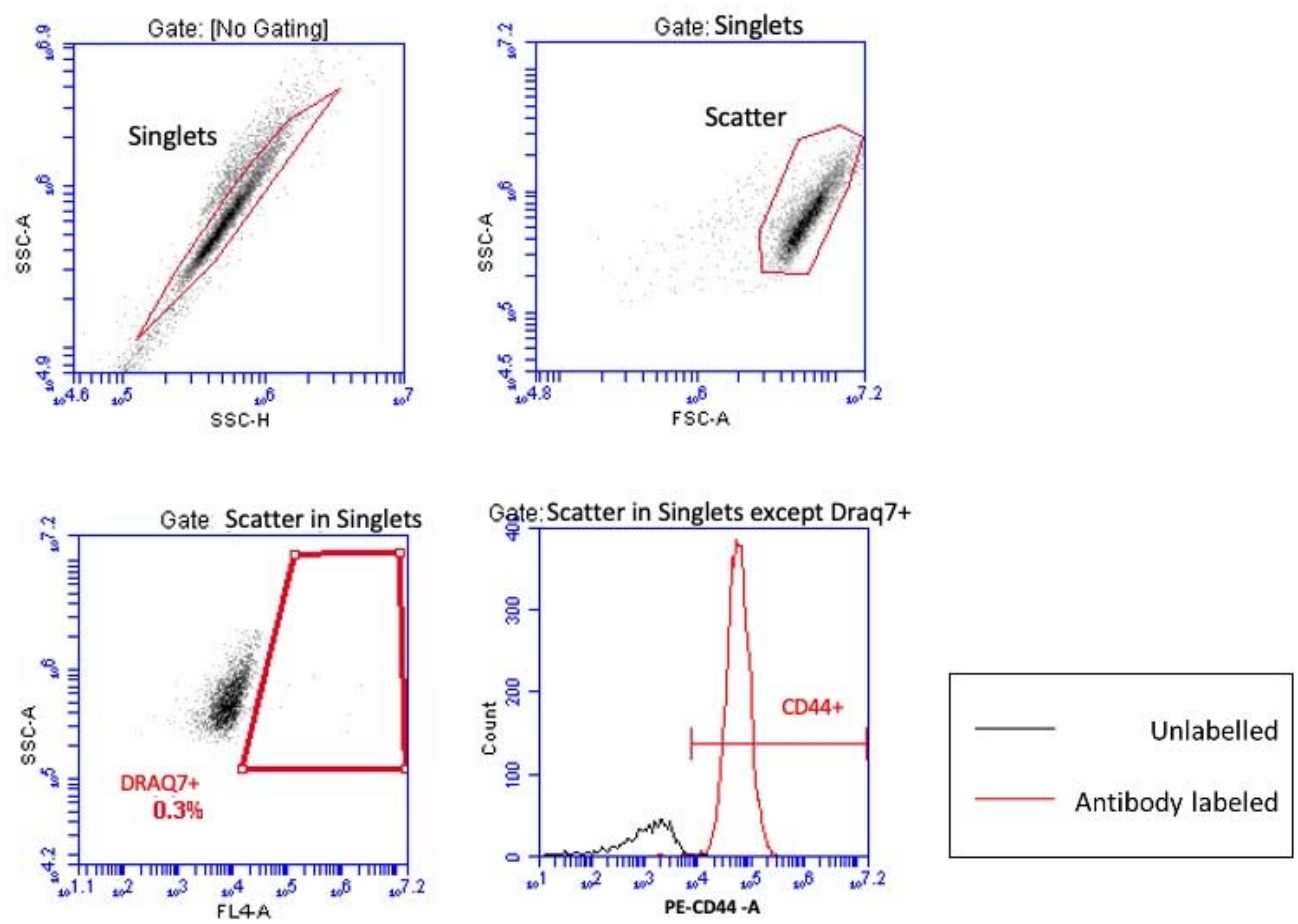

### Expected outcomes

When following the outlined protocol, both primary umbilical cord MSC and hTERT20-BMMSC (an immortalized bone marrow MSC cell line ^18, 19^kindly provided by Prof. Kassem, University Hospital of Odense, Odense, Denmark) were successfully used to form spheroids. MSC were seeded at densities of 15,000 cells per well. After 3 days of culture, visible spheroids formed in all wells. After culturing for 3 days, spheroid dissociation was performed as per the described method. Two dissociation cycles were needed for dissociate single cells from the spheroids formed. As shown in Figure 2, cells dissociated from the spheroids retained MSC surface marker expression (CD73 and CD44) and did not express hematopoietic markers (CD34 and CD45). Cells isolated from spheroids showed high viability (87%) despite accutase treatment.

hTERT20-BMMSC spheroids were differentiated into endothelial-like cells over 11 days following 3 days of initial spheroid formation with 15,000 cells per well. As illustrated in Figure 1, the endothelial markers vWF and CD31 were expressed in the differentiated spheroids. Notably, CD31 expression was not observed in previous 2D MSC endothelial differentiation assays conducted in our lab, even after 23 days of treatment with endothelial differentiation medium.

**Figure 1.**
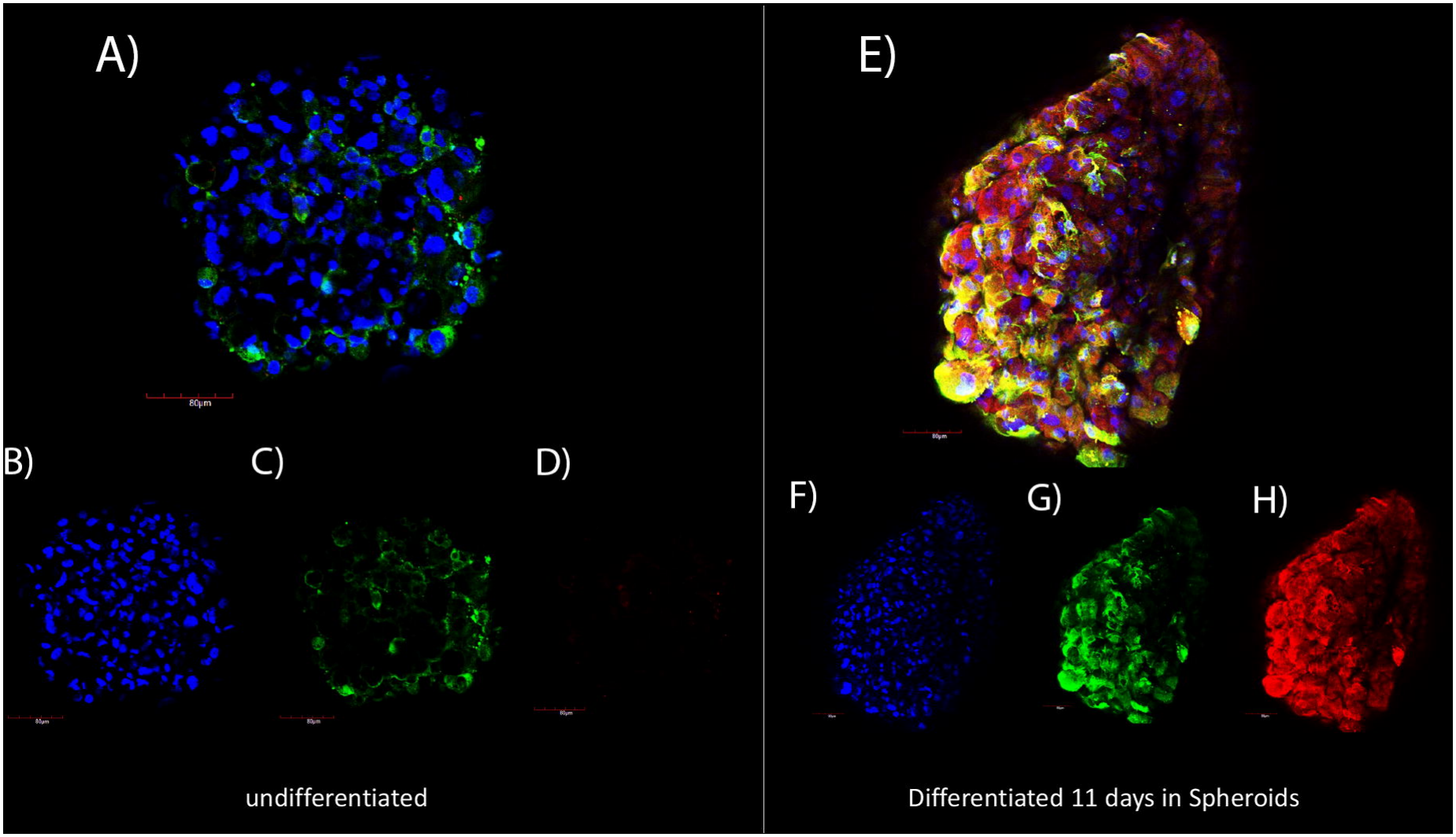
Immunofluorescence images for undifferentiated (A-D) and differentiated (E-H) hTERT20-BMMSC spheroids. Images were taken with Olympus FV1000 confocal microscope. One representative result is shown from 3 independent experiments. A & E: overlapped 3 channels (DAPI, vWF and CD31), B& F: DAPI channel, C& G: vWF channel, D& H: CD31 channel. Blue: DAPI, Green: vWF, Red: CD31. Scale bar= 80 µm

**Figure 2.**
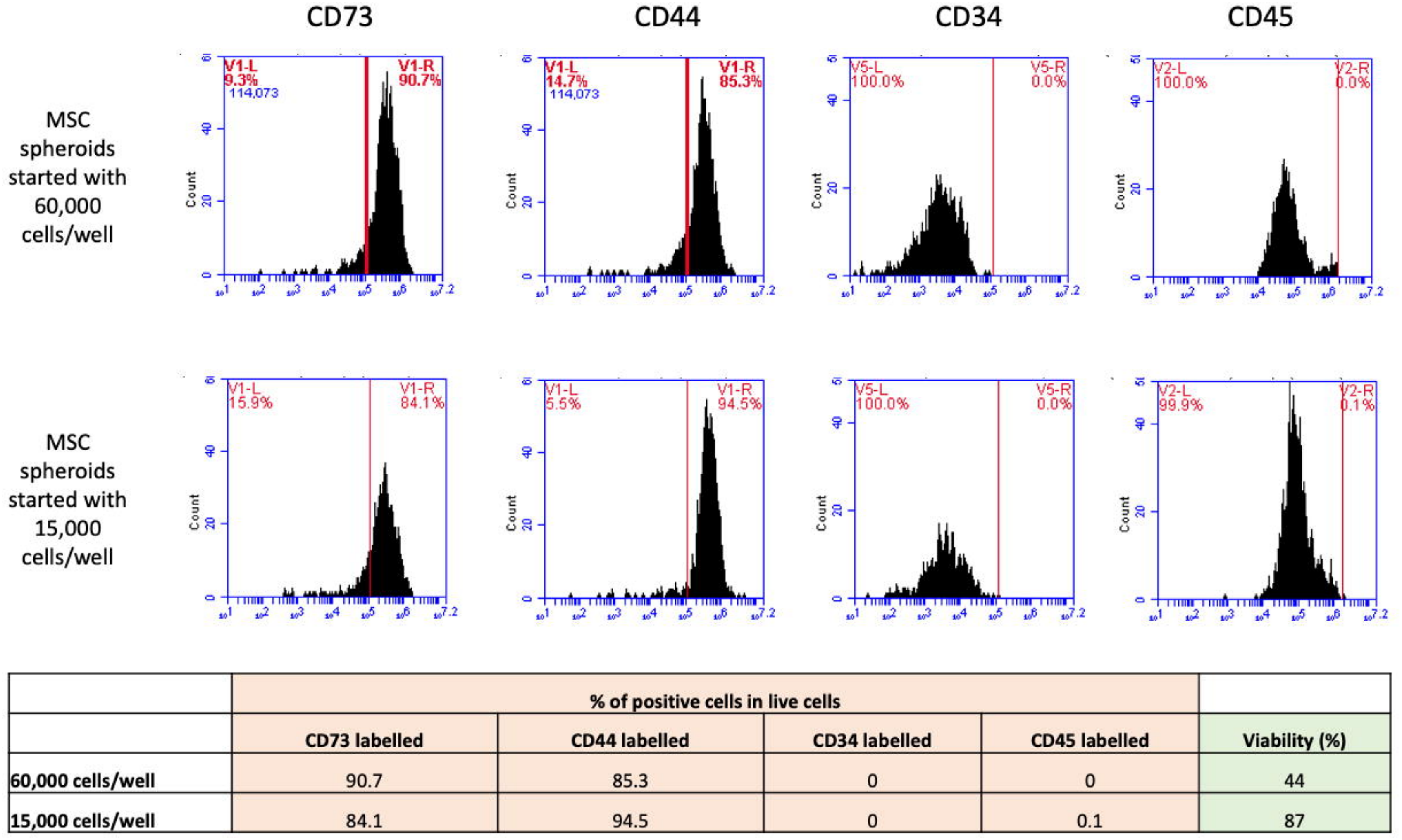
MSC surface markers expressed by MSC disassociated from hTERT20-BMMSC spheroids which were cultured for 3 days. Results were analysed by BD Accuri C6 flow cytometer. Analysing gates were set according to unlabeled negative control samples.

Dissociation of spheroids was performed on both endothelial-differentiated MSC spheroids, with undifferentiated MSC spheroids as a control group. Growth factors-induced differentiation was carried out over 11 days, following an initial 3-day spheroid formation period. As shown in Figure 3, CD44 expression in cells from both undifferentiated and differentiated spheroids showed a significant reduction, dropping to 83.71% and 73.15%, respectively, compared to over 90% in MSCs isolated from short-term cultured spheroids (3-day). One possible explanation involves the role of CD44 in MSC migration and adhesion. Qian et al. reported that primary MSCs isolated from bone marrow are initially CD44-negative but gradually acquire CD44 expression during in vitro culture^20^. In our method, MSCs are prevented from adhering to any extracellular matrix or culture surface, facilitating spheroid formation. This non-adherent culture environment may explain the reduced CD44 expression observed in MSCs dissociated from long-term cultured spheroids. This decrease in CD44 could also indicate the loss of MSC stemness and that the spheroid environment may enhance the differentiation potential of MSCs.

**Figure 3.**
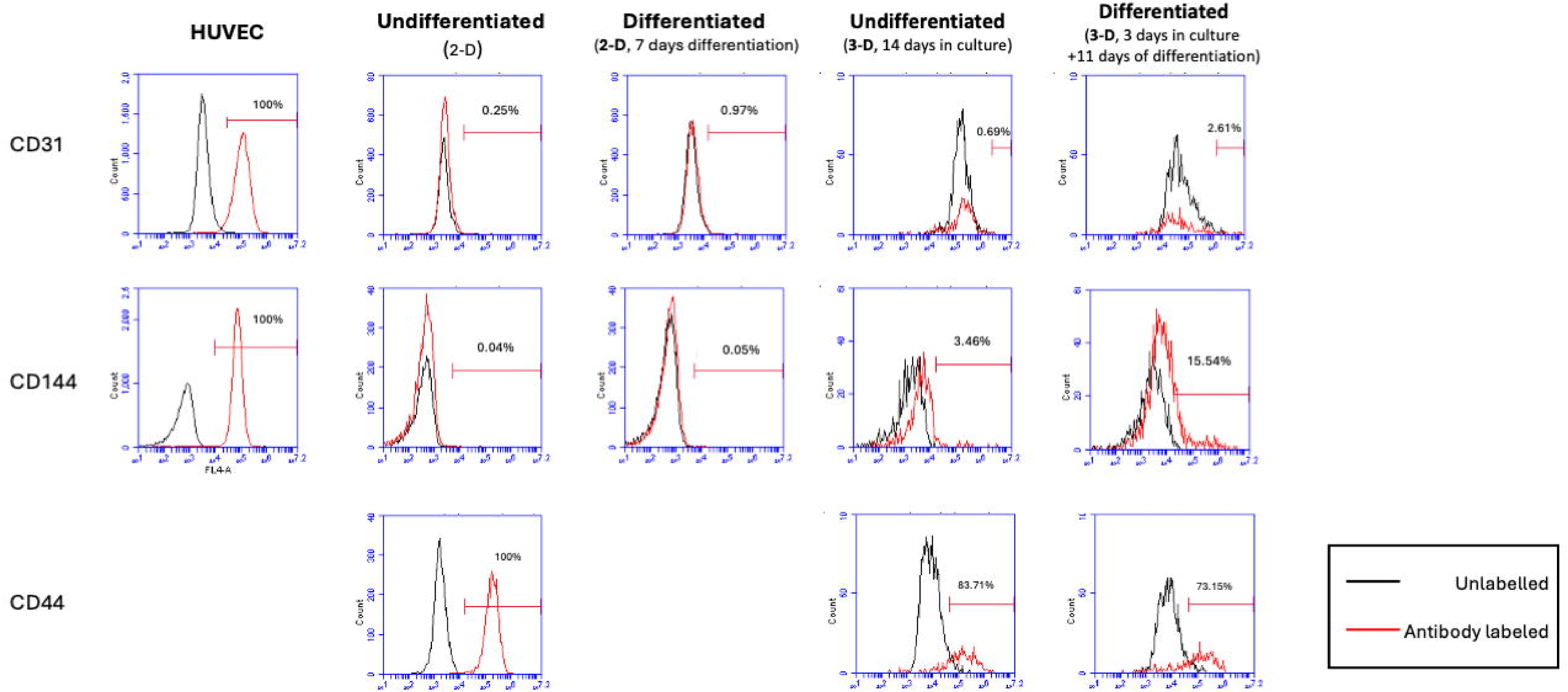
MSC surface marker CD44 and endothelial markers (CD31, CD144) expressed by MSC differentiated in 2-D culture and MSC disassociated from undifferentiated and differentiated spheroids. Results were analysed by BD Accuri C6 flow cytometer. Analysing gates were set according to unlabeled negative control samples. HUVEC: human umbilical vein endothelial cells. One representative experiment was shown here.

However, when analysing endothelial markers in cells dissociated from both undifferentiated and differentiated spheroids, we observed only modest increases in the percentage of CD31 (2.61%) and CD144 (15.54%) positive cells after differentiation, compared to undifferentiated spheroids (0.69% and 3.46%, respectively; Figure 3). Although endothelial marker expression was generally low, the increase remained modest compared to CD31 and CD144 expression levels observed in differentiated MSCs cultured in 2D (0.97% and 0.05%, respectively, Figure. 3). These results differ from the high CD31 expression observed in the immunofluorescence assays (Figure 1). This discrepancy may be attributed to the difference in detection methods; immunofluorescence involves fixation and permeabilization, which allows for the detection of intracellular markers, whereas flow cytometry only measures surface expression. Additionally, dissociating spheroids using accutase may lead to a reduction in the expression of cell surface markers, which could explain the low percentages of endothelial markers detected by flow cytometry. Lai et al. demonstrated that incubating cells with accutase for 10 or 30 minutes led to a significant reduction in the expression of surface Fas ligand, suggesting this enzymatic treatment may affect the integrity of surface proteins, making it challenging to accurately measure their expression in flow cytometry assays ^21^. In this context, immunofluorescence may offer a more sensitive method for detecting endothelial markers that are endogenously produced but not yet fully expressed on the cell surface.

### Limitations

A limitation of this study is that after long-term incubation, the exact number of MSC within each spheroid remains undetermined. This poses a challenge in achieving consistent outcomes when performing assays that rely on precise cell counts. The protocol was developed as part of a project aimed at utilizing MSC as a personalized source of endothelial cells to predict endothelial damage following allogeneic hematopoietic stem cell transplantation (HSCT). To accomplish this, it is critical to differentiate MSC into endothelial-like cells, with the degree of similarity between the differentiated MSC and authentic endothelial cells being a key factor in the predictive accuracy of the model.

Additionally, we observed that differentiated spheroids do not always maintain a perfectly round shape, potentially due to uneven differentiation rates across different parts of the spheroid. This could also be attributed to the heterogeneity of the MSC population used in this assay, where variable cell characteristics and differentiation potentials might contribute to the non-uniformity of the spheroid structure. Such heterogeneity in MSC populations could result in regions of the spheroid differentiating at different rates or displaying distinct characteristics, complicating downstream applications.

### Troubleshooting

#### Problem 1

Multiple spheroids can be formed in one well (related to step 7).

#### Potential solution

Seeding MSCs at a density of 60,000 cells per well often results in the formation of multiple spheroids. The presence of multiple spheroids suggests that the seeding density is too high or that cell viability is suboptimal. To address this issue, ensure that cell viability exceeds 95% prior to seeding for spheroid formation. Additionally, the seeding density may need adjustment, particularly when using different MSC lines or primary MSCs, as they may exhibit distinct spheroid-forming behavior. A lower seeding density may be required to achieve optimal spheroid formation with other MSC sources.

#### Problem 2

Fluorescence signals representing markers assessed by immunofluorescence are notably weaker in the core of the spheroids. This may be due to insufficient fixation and/or permeabilization (related to steps 12 and 14)

#### Potential solution

The duration of fixation can be adjusted based on spheroid size. In our studies, spheroid diameters ranged from 430 to 540 µm, with differentiated spheroids being larger than undifferentiated ones. For spheroids of this size, a fixation time of one hour is adequate. For larger spheroids or more robust fixation, the duration of step 12 and 14 can be extended, and the process can be performed on a rocker to ensure thorough penetration.

#### Problem 3

DAPI signal is weaker in the core of the spheroids. This could be related to insufficient staining duration of DAPI (related to step 21) or insufficient fixation/permeabilization (discussed in Problem 2.)

#### Potential solution

The duration of DAPI staining can be adjusted if the signal intensity is weak in the spheroid core. Alternatively, the staining step can be performed on a mixer with temperature control set to 37°C, which may enhance staining penetration and overall signal quality.

#### Problem 4

Difficulty when disassociating cells from spheroids. (related to Step 30)

#### Potential solution

Insufficient dissociation cycles may result in incomplete cell recovery from the spheroids. Increasing the number of dissociation cycles can improve dissociation efficiency. If four cycles are inadequate, the spheroids may be too large, indicating a need to adjust the seeding density. Alternatively, the dissociated cells can be collected and transferred to culture medium, allowing additional dissociation cycles to be applied to the remaining spheroids without further damage to the dissociated cells.

#### Problem 5

Low viability of cells observed after spheroids dissociation

#### Potential solution

Cell viability can be assessed before or during flow cytometry analysis. While dissociation may reduce cell viability, it should ideally remain above 70%. Low viability may result from excessive Accutase treatment or insufficient nutrient supply. To mitigate this, consider replacing the medium daily rather than every 2–3 days to ensure adequate nutrient availability.

## Resource availability

### Lead contact

Further information and requests for resources and reagents should be directed to and will be fulfilled by the lead contact, Guenther Eissner.

### Technical contact

Technical questions on executing this protocol should be directed to and will be answered by the technical contact, Simeng Li.

### Materials availability statement

This study did not generate new unique reagents.

### Data and code availability statement

All relevant data are included within the manuscript and its Supporting Information files. As this manuscript does not include any statistical analysis or graphical data, there are no additional datasets to share. The raw flow cytometry data (FCS files) are included as supporting information.

## Acknowledgments

The funding used for this study is provided by National Children’s Research Centre (NCRC), Ireland (funding no. A-18-4). We would like to acknowledge the help and support of Dr Alfonso Fernández, Prof. Jeremy Simpson and Dr Dimitri Scholz to this research protocol.

## Author contributions

**Conceptualization:** SL. **Methodology:** SL and IAQ. **Data Curation:** SL and IAQ. **Investigation:** SL and IAQ. **Resources:** GE and OS. **Writing – Original Draft:** SL. **Writing – Review & Editing:** GE and WK. **Supervision:** GE. **Project Administration:** GE and WK. **Funding Acquisition:** GE and OS. **Visualization:** SL. **Validation:** SL and IAQ.

The funders had no role in study design, data collection and analysis, decision to publish, or preparation of the manuscript.

## Declaration of generative AI and AI-assisted technologies in the writing process

During the preparation of this work the author(s) used ChatGPT in order to check grammars and proofread. After using this tool/service, the author(s) reviewed and edited the content as needed and take(s) full responsibility for the content of the publication.

